# Magnetic Particle Imaging Lymphography (MPIL): A novel technique for lymph node mapping

**DOI:** 10.1101/2025.06.11.658392

**Authors:** Nitara Fernando, Olivia C. Sehl, Benjamin Fellows, Chinmoy Saayujya, Paula J. Foster

## Abstract

During metastasis, tumour cells drain to nearby lymph nodes, the first of which are named the sentinel lymph node(s) (SLN). SLN biopsy (SLNB) determines if metastasis has occurred. Traditionally, SLNB involves injecting a Technetium-labeled tracer peritumourally, pre-operative imaging with SPECT to locate the SLN, and surgery guided by a gamma probe to remove them. Limitations include short tracer half-life which can make scheduling the SLNB difficult, and radiation dose to patients and healthcare workers. Alternatively, magnetic localization, with superparamagnetic iron oxide nanoparticles (SPIONs) as the tracer and a magnetometer to detect SPIONs in the SLN during surgery can be used, however, this lacks pre-operative imaging.

Magnetic Particle Imaging (MPI) is a new imaging modality that directly detects SPIONs, holding potential for pre-operative imaging in magnetic SLNB. SPIONs for SLNB should have rapid drainage, high SLN accumulation, and high specificity to the SLN. For MPI Lymphography (MPIL), high particle sensitivity is also important. This study assesses the in vivo pharmacokinetics for SLN mapping with MPIL in a murine model, using five commercially available SPIONs of varying iron core sizes and surface coatings. We show that some SPIONs provide higher MPI signal at the SLN and show the potential to detect higher echelon nodes (HENs). PEGylation of SPIONs and mannose targeted SPIONs increase clearance from the injection site and the mannose targeted SPION reduces flow to HENs.

## Introduction

### Overview of Sentinel Lymph Node Biopsy (SLNB)

Cancer remains a major health problem worldwide, with metastatic disease being responsible for most tumour-related deaths.^1^ Many types of cancer metastasize through the lymphatic system, and one of the earliest sites of spread for these cancers are nearby lymph nodes.^2,3^ The sentinel lymph node (SLN) is the first lymph node (or group of nodes) to which cancer cells are most likely to spread from a primary tumour. To determine if metastasis has occurred, the SLN is identified, removed, and examined by pathology to determine if cancer cells are present.

Sentinel lymph node biopsy (SLNB) is the primary method of surgical management of the regional lymph node basin and staging for most breast cancer and melanoma patients, and this procedure is expanding to new indications such as head and neck cancer (HNC) and gynecological cancers.^4–7^ The common clinical workflow consists of a peritumoral, subdermal injection of a radioactive technetium-based nanocolloid which quickly drains to, and accumulates in, the SLN. This is often combined with a blue dye injection to improve visualization of the lymphatic tract intraoperatively. Pre-surgical imaging using scintigraphy or SPECT is then performed to identify the location of the SLN(s) which is marked on the skin. Next, an incision is made and intraoperative mapping is performed using a gamma probe to identify nodes with high radioactive counts which are removed and then histologically assessed for cancer cells.^4–7^ In order to determine which nodes are removed, a ‘10% rule’ is commonly used intraoperatively, where nodes with the highest radioactive counts as well as those that are 10% or more of the ex vivo count of the hottest node are removed.^8,9^ This rule is used to define radioactive nodes as either relevant SLNs that are more likely to contain metastasis, or irrelevant higher echelon nodes (HENs) that do not need to be removed.

There are several drawbacks of the nuclear medicine SLN localization procedure. Logistically, the main challenge is the short (6 hour) half-life of the tracer, which can cause challenges in scheduling SLN mapping prior to surgery.^10^ The short half-life also means the radiotracer must continually be produced, a long-term supply cannot be stockpiled, and tracer shortages are common. Furthermore, many hospitals in the world do not have access to radioisotopes.^11,12^ There is also radiation dose to patients and health care workers, and the requirement for special handling and specific regulatory requirements. A “shine-through” phenomenon has also been reported, where radioactive signal at the injection site produces a large hotspot which can hide SLNs.^13^ The cutoff point for radioactivity, and the 10% rule, have been scrutinized for possibly contributing to the excessive removal of negative nodes.^14^

### Magnetic SLNB

Magnetic SLNB is becoming a viable alternative to the standard nuclear medicine technique for breast, prostate, oral, and gynecological cancers.^15–21^ Like conventional SLNB, a tracer is administered peritumorally and drains quickly to the SLNs. This tracer is a superparamagnetic iron oxide nanoparticle (SPION), such as Resovist, Magtrace, Sienna+, or Ferrotrace. A hand-held magnetometer is used intraoperatively (like the gamma probe) for magnetic mapping to detect nodes with magnetic signals, guiding the SLN dissection. The same cutoff used for the gamma probe has been applied for magnetic SLNB, in this case any node with a magnetometer count of 10% or more of the node with the highest count is excised. The SPIONs have the added benefit of providing a color change in the node (brown or black) and extensive tracer half-life (> 9 days), which provides flexibility for surgical planning.^15^

The magnetic SLN localization technique also has limitations. The transcutaneous magnetic signal is weak, so it does not provide pre-operative information on SLN location, therefore, it is mainly used for breast cancer which has predictable drainage patterns.^22^ For other cancers with complex anatomies (i.e., HNC) or less predictable drainage (i.e., melanoma of trunk and extremities) presurgical imaging is fundamental as SLNs can be located at several, and even unpredictable, locations. Lingering skin discolouration caused by high doses of SPIONs has also been reported.^23^

### Magnetic Particle Imaging (MPI)

Magnetic Particle Imaging (MPI) is an emerging imaging technology that directly detects SPIONs (the same SPIONs used for the magnetic SLNB). The MPI signal is only generated when the magnetic moments of the SPIONs rotate in response to applied fields, so the signal is specific to iron. Biological tissues neither generate nor attenuate MPI signals. This imbues MPI with a positive “hotspot” contrast (like SPECT and PET) that provides spatial localization and quantification without ambiguity. The MPI signal is linearly quantitative with SPION concentration, and therefore the amount of iron can be directly calculated. A study by Sehl et al., showed the feasibility of MPI to sensitively detect and quantify SPION signal in lymph nodes using various murine lymphatic drainage models.^24^ The purpose of this study was to compare various SPIONs for MPI lymphography (MPIL).

## Methods and materials

### Magnetic particles

Five SPIONs were compared in this study, as outlined in **Table 1**. These SPIONs were chosen to compare the effects of iron core size and particle coatings. VivoTrax (Magnetic Insight Inc.) is a hydrophilic colloidal solution of SPIO coated with carboxydextran. The iron core size is bimodally distributed— reported to have approximately 70% of the solution consisting of ∼5 nm cores, and 30% consisting of ∼24 nm cores. The mean hydrodynamic diameter is 62 nm. VivoTrax+ is magnetically fractionated to select for larger iron cores, 70% 24 nm cores, and 30% 5 nm cores. The hydrodynamic diameter is 62 ± 4 nm.

**Table 1.**
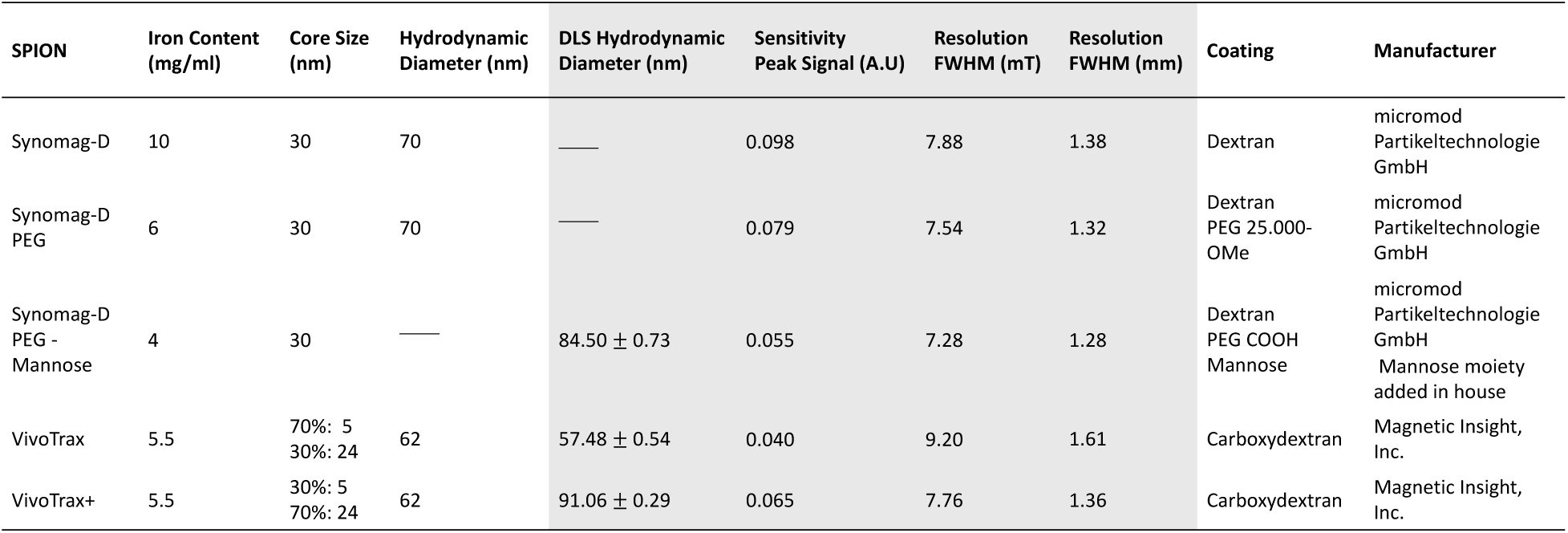
Comparison of physical and magnetic properties of commercial SPIONs. Manufacturer provided information including iron content, core size, hydrodynamic diameter, and coating is included. DLS measured hydrodynamic diameter is provided. MPI relaxometry measured sensitivity (A.U), and resolution (mT) are provided. A magnetic gradient of 5.7 T/m is used to approximate resolution in mm. The grey shaded regions of the table represent in house acquired data.

**Table 2.** Iron (Fe) in SLNs estimated using MPI and ICP-MS. MPI signal on day 8 is used to estimate the Fe in the SLN on day 8 using an injected dose of 0.8 mg/kg. The range (minimum-maximum) of Fe injected, in vivo MPI signal on Day 8, and Fe on day 8 for n = 4 mice from each SPION is shown. For one mouse from each SPION, ICP-MS was performed on the ipsilateral and contralateral popliteal nodes to determine Fe content. ^++^Vivotrax+ estimated Fe from MPI is based on the 2 cm FOV quantification, while for other SPIONs it is from 12 cm quantification.

|  | MPI (n=4) |  |  | ICP-MS (n=1) |  |
| --- | --- | --- | --- | --- | --- |
| | Fe injected<br>[$\mu\text{g}$] | MPI Signal<br>Day 8 [%ID] | Fe on Day<br>8 [$\mu\text{g}$] | Fe in Ipsilateral<br>SLN [$\mu\text{g}$] | Fe in Contralateral SLN<br>(control) [$\mu\text{g}$] |
| Synomag-D Plain | 18.4 – 20.8 | 14.53 – 24.47 | 2.96 – 5.04 | 4.35 | 1.16 |
| Synomag-D PEG | 16.8 – 23.3 | 19.64 – 23.52 | 3.30 – 4.78 | 2.49 | 0.83 |
| Synomag-D PEG-Mannose | 21.6 – 26.4 | 7.12 – 18.90 | 1.89 – 4.54 | 1.16 | 0.58 |
| VivoTrax | 15.2 – 17.6 | 2.86 – 5.61 | 0.44 – 0.99 | 1.42 | 0.69 |
| **VivoTrax+ | 15.2 – 18.4 | 1.97 – 5.28 | 0.30 - 0.93 | 0.87 | 0.96 |

Synomag-D particles (Micromod GmBH) are multicore particles with a nanoflower substructure.^25^ The agglomerated iron core size is ∼30 nm and is coated with dextran to improve stability and provide biocompatibility. The hydrodynamic diameter is ∼70 nm. We compared dextran-coated Synomag-D (Plain Synomag-D) with two alternate coatings: (1) Polyethylene glycol (PEG) and (2) PEG-Mannose. A PEG-coated version of Synomag-D (Synomag-D PEG 25.000-OMe) is available through the manufacturer and has the same hydrodynamic diameter (70 nm) and iron core size (∼30 nm) as the Plain Synomag-D.

Coupling of D-Mannosamine to the carboxyl functional Synomag-D-PEG-COOH particles, was done using simple carbodiimide chemistry. All solutions were made up in 0.5 M 2-(N-morpholino)ethanesulfonic acid buffer unless stated otherwise. Briefly 2.5 mL of 10 mg/mL of COOH functional magnetic nanoparticles were combined with 100 μL of a 40 mg/mL 1-Ethyl-3-(3-dimethylaminopropyl)carbodiimide solution and 50 μL of a 30 mg/mL N-hydroxysulfosuccinimide (sulfo-NHS) solution. This combined solution was left to react on a shaker table for 45 minutes. After 45 minutes 1 mL of 150 mg/mL mannosamine-HCL was added to the reaction and left to shake for another 3 hours to complete the coupling. After 3 hours, the combined solution was transferred to 10 k MW cutoff dialysis tubing and dialyzed against 0.01 M phosphate-buffered saline (PBS) for 24 hours. The total volume of the particle solution remaining in the dialysis tubing was measured and particle concentration of the final solution was calculated based on the 25 mg of particles added at the beginning of the reaction.

### Dynamic Light Scattering (DLS)

Dilute nanoparticle samples were subjected to DLS using a Brookhaven NanoBrook Omni Light Scattering System (Holtsville, NY) to determine the distribution of hydrodynamic diameters. Three independent measurements were made for Vivotrax, Vivotrax+, and Synomag-D PEG-Mannose, then average ± standard deviation values are reported.

### Magnetic particle relaxometry (MPR)

Magnetic particle relaxometry (MPR) was performed on triplicate samples for each SPION on the Momentum™ scanner. MPR is a method to characterize SPIONs by measuring their net magnetization and relaxation by turning off the selection field and applying a sweeping magnetic field (negative to positive) back and forth. This causes the SPIONs to go from negative to positive magnetic saturation, creating a point spread function (PSF) as the output. The PSF output is used to measure particle resolution by measuring the full width half maximum (FWHM), and sensitivity, by measuring at the peak height. PSFs were analyzed using Prism Software (Version 10.3.1).

### In vivo imaging

Healthy, nu/nu mice (n = 20) were purchased from Charles River Laboratories, Inc. (Senneville, CAN). All animal studies were performed in accordance with institutional and national guidelines. First, mice were anesthetized with isoflurane administered at 2% oxygen. Subsequently they were administered 0.8 mg/kg of SPION, in a 40 uL PBS solution (n = 4 for each; **Table 1**), intradermally to the left footpad. This injection route models an established drainage pathway from the footpad to the popliteal lymph node (pLN), defined as the SLN in this model.^26^ Although “SLN” is a term typically used in the context of cancer, these experiments use healthy mice. This enables characterization of normal lymphatic drainage pathways and facilitates the evaluation of this novel SLN mapping technique without the confounding influence of tumour-associated lymphatic alterations.^24,27^

All imaging was performed using the Momentum™ pre-clinical MPI scanner (Magnetic Insight Inc.). The Momentum^TM^ is an oil-cooled, field free line (FFL) scanner that reconstructs images using x-space signal processing.^28^ 2D tomographic images were acquired with a 5.7 T/m selection field gradient. At each timepoint, full body field of view (FOV) (6 x 6 x 12 cm (X, Y, Z)), multichannel (drive field strengths: x = 13 mT, z = 20 mT, scan time ∼ 2.5 minutes) scans with the native reconstruction (inverse combiner = off) were acquired. Subsequently, a focused 2 cm FOV (6 x 6 x 2 cm (X, Y, Z)) was prescribed for the acquisition of single channel scans (drive field strength: z = 26 mT, scan time ∼ 2.5 minutes) which were reconstructed with a tailored reconstruction algorithm available in the momentum advanced user interface (inverse combiner = on) centered around the SLN. For one mouse from each SPION, 3D scans were acquired at the t = 4 hour timepoint using a full body FOV, multichannel (drive field strengths: x = 13 mT, z = 20 mT, 35 projections, scan time ∼ 35 minutes). We have previously described and tested these image reconstruction methods and found that using these parameters allows for optimized small FOV images that are quantifiable.^29^ Mice were imaged with MPI 10-20 minutes post-injection, to determine a baseline signal, and then 4, 24, and 192 hours post-injection to quantify signal to the SLN. Mice were fasted 12 hours prior to each imaging timepoint to minimize gut signal due to iron in mouse feed (**Figure S1A**).

Magnetic Resonance Imaging (MRI) was performed for one mouse administered VivoTrax, t = 4 hours after injection. MRI was acquired on a 3T clinical scanner (Discovery MR750, General Electric) using a 4.31 x 4.31 cm diameter RF surface coil (Clinical MR Solutions, Wisconsin). An iron-sensitive balanced steady state free procession (bSSFP) imaging sequence was used, with the following imaging parameters: TR/TE = 12.8/6.4 ms, flip angle = 20°, field of view = 6.4 cm x 3.2 cm x 2.6 cm, resolution = 400 µm isotropic, phase cycles = 12, bandwidth = ± 31.25 kHz, acquisition time = ∼10 minutes.

### Ex vivo imaging

Immediately following the last MPI exam, mice were sacrificed. Contralateral (control) and ipsilateral SLNs and HENs were removed from all mice and placed in 4% paraformaldehyde for 24 hours for fixation. Then, lymph nodes were transferred to a 70% ethanol solution. 2D MPI images were acquired from individual pLNs and HENs using the same parameters used for in vivo imaging (native reconstruction, 12 cm FOV, multichannel).

### MPI analysis and quantification

All MPI images were imported into Horos™, an open-source clinically relevant image analysis software (version 4.0.0, Annapolis, MD USA). Images were viewed using a custom MPI colour look-up table (CLUT).

For analysis of SLN signal, a threshold value of 0.5 times the maximum lymph node signal was applied as a lower bound for signal detection. This method was chosen as it allowed us to ensure that no signal from the injection site or surrounding lymph nodes were included when quantifying signal in a specific lymph node. Using this threshold, a semi-automatic segmentation tool was used to measure the mean MPI signal within a specific region of interest (ROI). Total MPI signal for an ROI was calculated by multiplying the ROI area (2D) by the mean signal. All MPI images were segmented and analyzed in the same way to ensure consistency.

To compare MPI signal at the SLN over time for all the tested SPIONs, percent injected dose (%ID) was calculated using **Equation 1**. Baseline signal was the summation of total MPI signal at the first imaging timepoint (10-20 minutes) and included signal at the injection site and signal at the SLN, if present.

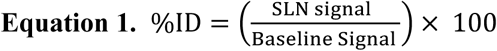

For analysis of HEN signal, a custom circular ROI of the same size (22.95 mm^2^) was drawn around each node (pLN, sciatic, iliac, renal) as shown in **Figure S1B**. This method was chosen, as opposed to the method used for SLN quantification, due to the proximity of HENs to the SLN, making it difficult to isolate signals. Total MPI signal for an ROI was calculated by multiplying the ROI area (2D) by the mean signal.

MPI signal at HENs was quantified as percent of SLN signal (%SLN) using **Equation 2**. This was done to relate HEN signal to the clinically relevant 10% rule.

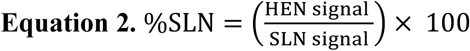

To analyze the signal at the injection site (footpad), a custom circular ROI of fixed size (111.6 mm²) was drawn. This approach was chosen to minimize the impact of signal spread at the injection site over time. Using a ROI of a fixed size allowed us to maintain a consistent ROI area, enabling more accurate quantification.

### Histopathology

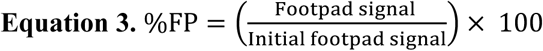

Following ex vivo imaging, excised contralateral (control) and ipsilateral SLNs, and HENs were paraffin embedded, and sectioned (10 μm thickness). The entire lymph node was sliced, with ∼5 slides per node with 10-15 sections per slide. All slides were subsequently stained with Perls’ Prussian Blue (PPB) which uses a mixture of hydrochloric acid and potassium ferrocyanide to stain for iron (blue). Microscopy was performed using the Echo 4 Revolve Microscope (CA, USA). The slides were visually assessed for presence of iron staining.

### Acid digestion of lymph node tissue and Inductively Coupled Plasma Mass Spectrometry (ICP-MS)

One SLN from mice injected with each SPION was digested. Lymph node tissue was placed in 5 mL of 68-70% nitric acid which was heated to 90°C and left to evaporate to dryness yielding an oily yellow residue. 5 mL of 30% peroxide solution was added and heated to 90°C. Once the volume of the peroxide reduced to ∼200 µL, another 5 mL of 68-70% nitric acid was added and left at 90°C to evaporate to dryness. This final boil off resulted in a white/yellow salt left on the surface of the vessel which was dissolved in 10 mL of 2% nitric acid and filtered through a 0.22 µm nylon filter. Solutions were then tested by ICP-MS following standard procedures at the Biotron Research Center at the University of Western Ontario.

### Statistical Analysis

Statistical analyses were performed using GraphPad Prism Software. Paired t-tests were used to compare MPR results between different SPIONs. Two-way Analysis of Variance (ANOVA) tests with multiple comparisons were used to compare in vivo SLN signal between different SPIONs. p<0.05 was considered a significant finding.

## Results

### Magnetic Particle Relaxometry (MPR) Sensitivity

Figure 1 shows the peak MPI signal measured by MPR for the various SPIONs. These values are also listed in **Table 1**. Plain Synomag-D had the highest peak MPI signal, 2.5x higher than VivoTrax which has been the most used SPION for pre-clinical MPI. The peak signal for VivoTrax+ was 1.6x greater than that for VivoTrax, consistent with previous literature.^30^ The peak signal for the Synomag-D particles decreased from Plain dextran-coated, to PEG-coated, to PEG-Mannose-coated. The values for FWHM measured by MPR are shown in **Figure S2.**

**Figure 1.**
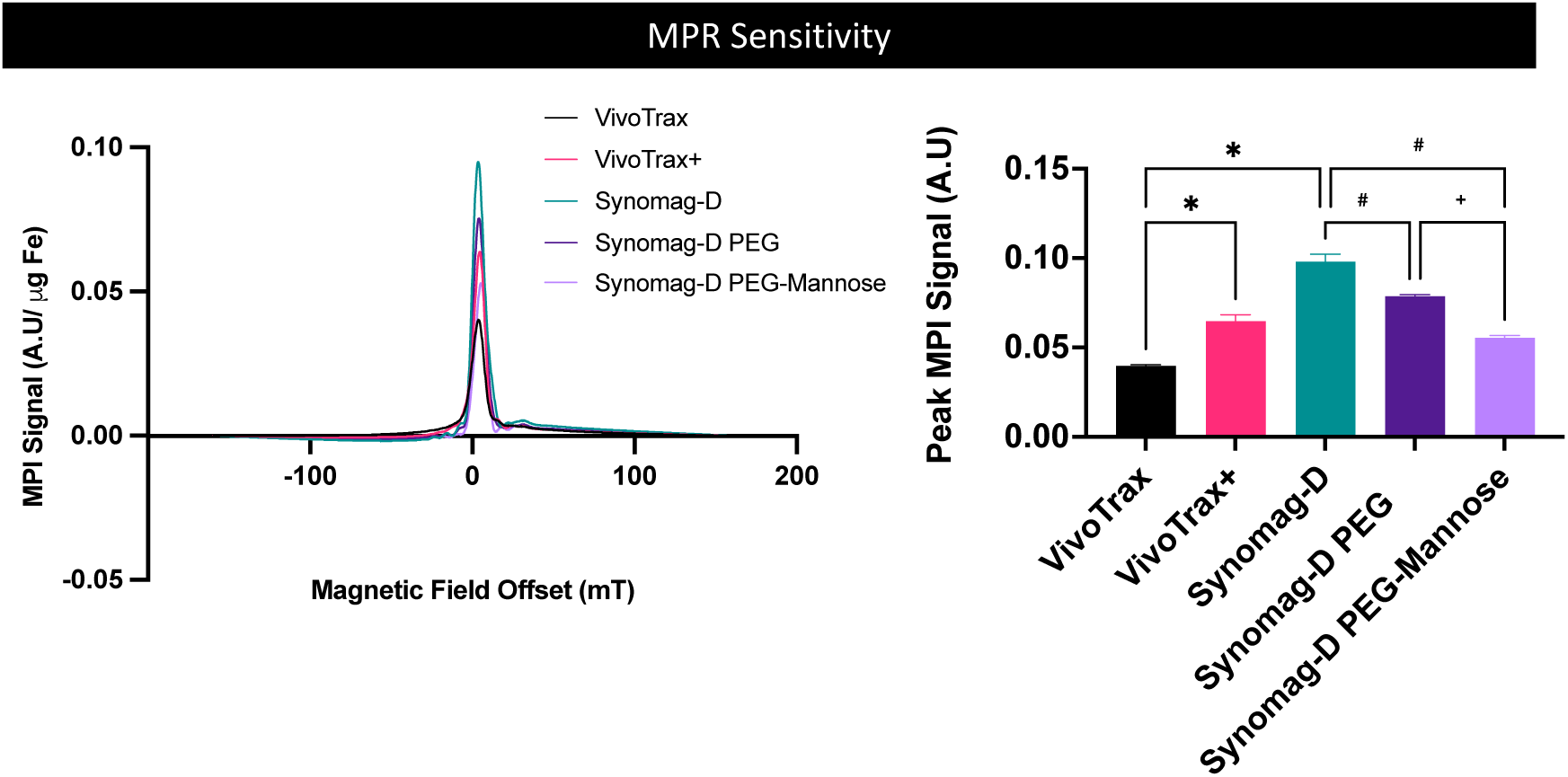
MPR Sensitivity. (A) Particle sensitivity measured by the peak height of the PSF is shown for all 5 SPIONs (n = 3 per SPION). The bar graph depicts the peak MPI signal for each SPION. (*p<0.05 (paired t-test with VivoTrax), ^#^p<0.05 (paired t-test with Synomag-D), ^+^p<0.05 (paired test with Synomag-D PEG)).

### In Vivo Imaging Comparing SPIONs of varying cores

Representative 2D full FOV mouse images acquired 4 hours after mice were injected with VivoTrax, VivoTrax+ or Synomag-D are shown in Figure 2A. MPI images are scaled to display the maximum MPI signal at the SLN. At 4 hours post-injection MPI signal was detected at the SLN for all 4 mice injected with Synomag-D or VivoTrax. For Synomag-D, MPI signal at the SLN was visible as early as 10-20 minutes post-injection and up to 8 days. For VivoTrax, at later timepoints (24 hours and 8 days) signal was only detected in 3 of 4 mice. No MPI signal was detected at the SLN for any of the mice administered VivoTrax+ at any timepoint. The percentage of the total injected dose at the SLN (%ID) was significantly higher at all timepoints for mice administered Synomag-D compared to mice administered the same dose of VivoTrax (Figure 2B). There was no significant difference in the %ID over time (between 4 hours and 8 days) for either Synomag-D or VivoTrax. For one mouse administered VivoTrax, MRI was also acquired 4 hours after injection (**Figure S3**). The signal void in the ipsilateral SLN showed good colocalization with the signal hotspot seen in MPI (**Figure S3C**).

**Figure 2.**
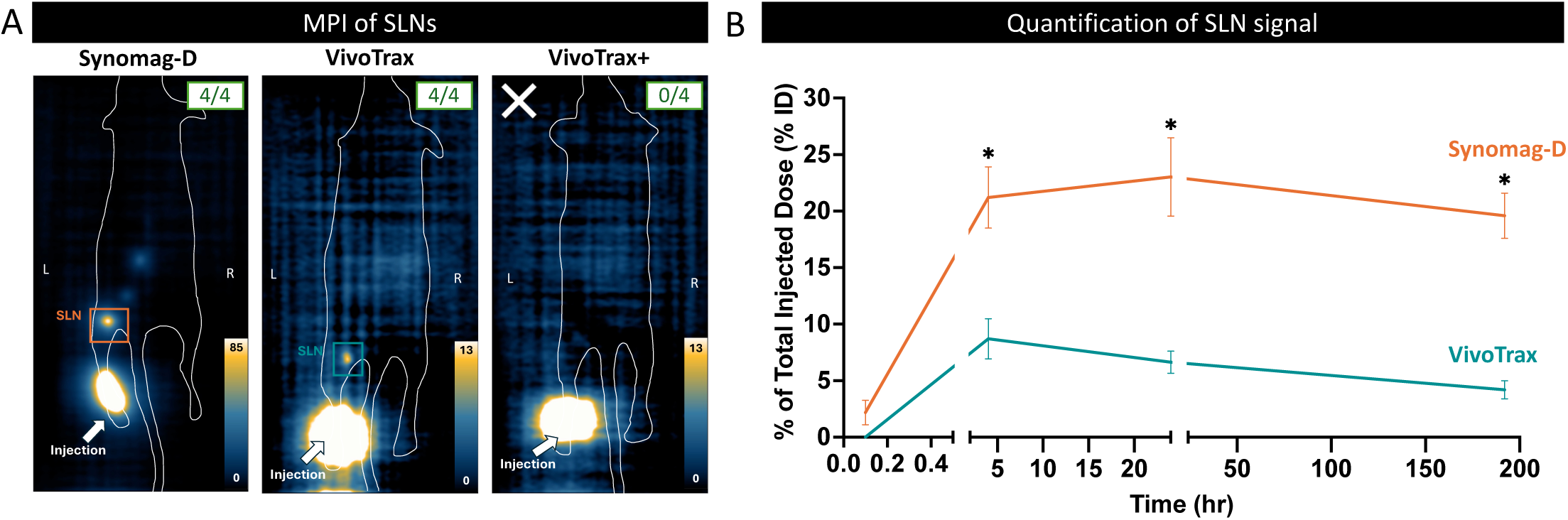
Comparing SPIONs of varying cores. (A) MPI shows in vivo pharmacokinetics for three nanoparticles, Synomag-D, VivoTrax, and VivoTrax+ after intradermal injection. A representative 2D MPI for one mouse from each SPION group acquired at 4 hours post injection is shown (n = 4 per group). The arrow denotes the injection site (left footpad), and a box is drawn around the location of the SLN. For VivoTrax+ the SLN was not detected in full body scans, denoted by the white X. On the top right corner of each image, a box with a fraction denotes the number of mice for which MPI signal was detected at the 4 hour imaging time point. For Synomag-D, SLN signal was detected for 4/4 mice, for VivoTrax SLN signal was detected for 4/4 mice and for VivoTrax+ 0/4 mice had SLN signal. (B) Signal at the SLN for each SPION was quantified over time as %ID. Signal in the SLN was not quantifiable for any mice injected with VivoTrax+. %ID for mice administered Synomag-D was significantly higher than those administered VivoTrax 4 hours, 24 hours, and 8 days post-injection (*p<0.05).

A smaller focused FOV was tested to isolate the lower signal at the SLN from the high signal at the injection site. This approach has previously been used to improve detection of MPI signals of varying dynamic range (i.e., shine-through effects).^29,31–33^ This is shown in Figure 3. A small FOV centered around the SLN allowed for the detection of SLN signal from all mice at all timepoints, including those administered VivoTrax+ for which SLN signal could not be detected with a full FOV (Figure 2). The %ID was significantly higher for Synomag-D compared to both VivoTrax and VivoTrax+ at all timepoints. There was no significant difference in the %ID for VivoTrax and VivoTrax+.

**Figure 3.**
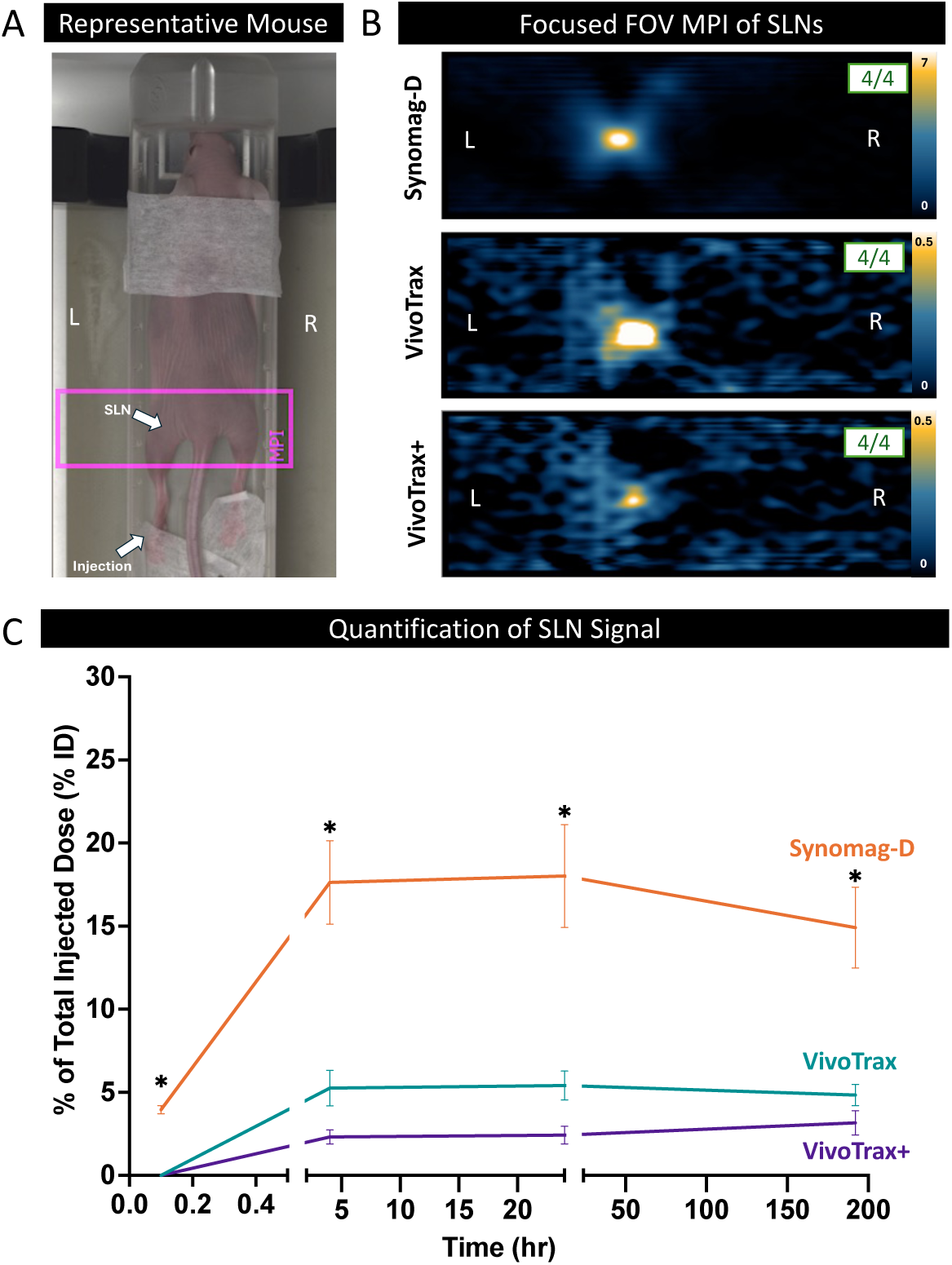
Focused FOV MPI of SPIONs. (A) A representative mouse on the imaging bed is shown. The pink box indicates where the focused FOV scan was positioned. The location is centered on the SLN. (B) Representative MPI of the focused FOV scans are shown for Synomag-D, VivoTrax, and VivoTrax+ with the signal scale bar on the right of each image indicating peak signal. On the top right corner of each image, a box with a fraction denotes how many mice MPI signal was detected at the 4 hours imaging time point. Signal can be detected using this scan mode for all SPIONs. (C) The signal is quantified as %ID over time for all three SPIONs. %ID for mice administered with Synomag-D was significantly higher than those injected with VivoTrax and VivoTrax+ at all timepoints (*p<0.05).

### Comparing SPIONs with Different Surface Coatings

The three Synomag-D particles were compared to assess the effects of different surface coatings on drainage to the SLN and HENs. Figure 4A shows representative 2D images acquired at the first (10-20 minutes) and last (8 days) timepoints. Plain Synomag-D had the fastest clearance from the injection site. At the first imaging timepoint, signal was detected at the SLN in all mice administered the Plain Synomag-D, in one mouse administered PEG-coated Synomag-D, and in none of the mice administered the PEG-Mannose-coated Synomag-D. MPI signal was detected at the SLN at all other timepoints in all mice (4/4) for all three Synomag-D particles. The quantification of %ID showed no significant differences in SLN signal between any of the Synomag-D particles, except for at 4 hours post injection where the signal for PEG-Mannose was significantly lower than PEG (Figure 4B). Signal at the injection site was quantified over time as %FP (Figure 4C). Plain Synomag-D had the lowest total clearance from the injection site. Between the first and last imaging timepoints the MPI signal in the footpad decreased by 11% for Plain Synomag-D, by 45% for PEG and by 49% for PEG-Mannose.

**Figure 4.**
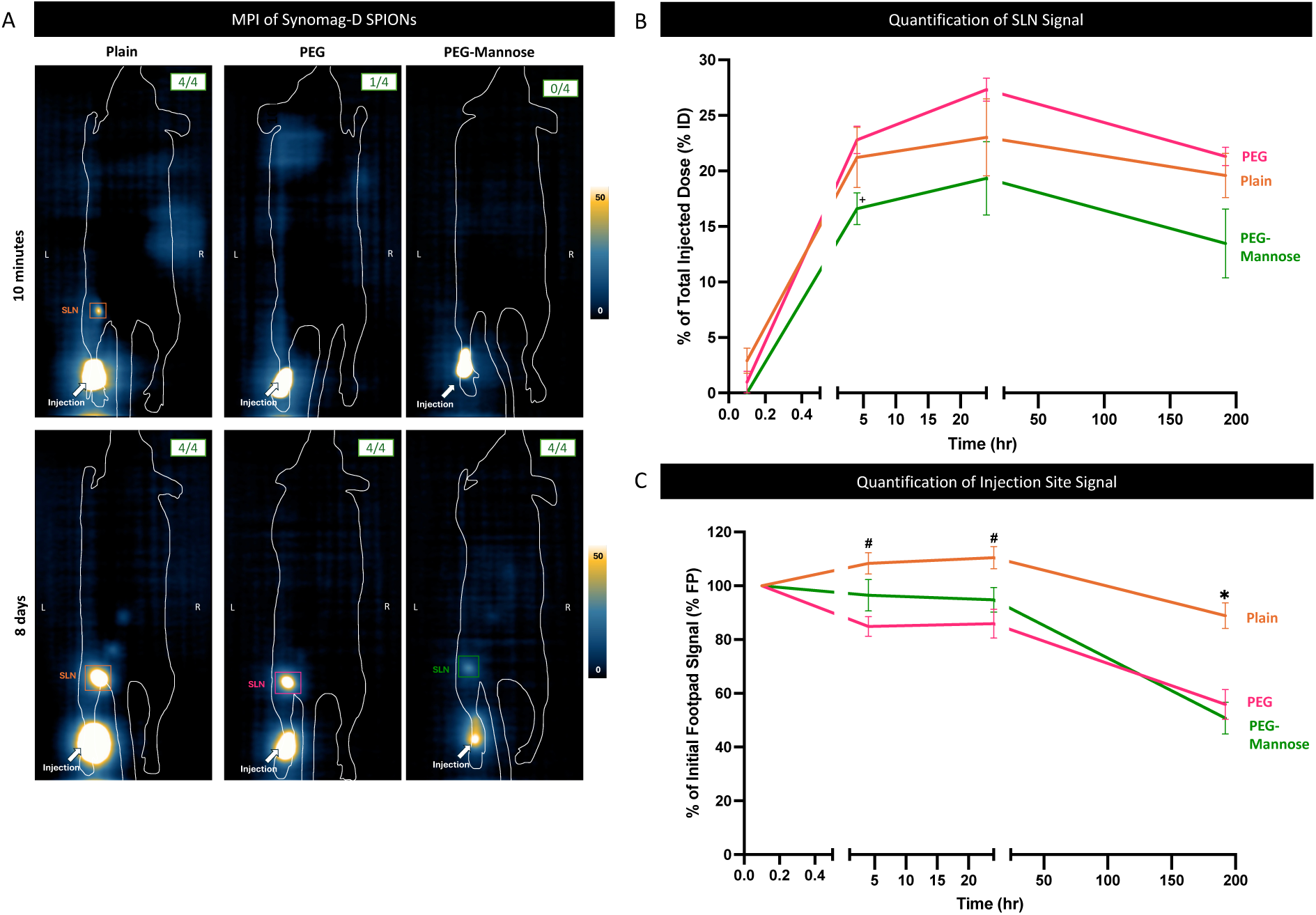
Comparing SLN Signal for Different Surface Coatings. (A) Representative 2D full FOV images are shown for mice administered Synomag-D with different surface coatings (Plain, PEG, and PEG-Mannose) 10-20 minutes after injection and 8 days after injection. (B) Signal to the SLN is quantified as %ID over time, there was no significant difference in SLN signal for the different surface coatings except for at t = 4 hours, where PEG and PEG-Mannose were significantly different (^+^p<0.05). (C) Signal at the injection site (footpad) is quantified as %FP over time, at t = 4 hours and t = 24 hours Plain Synomag-D is significantly different from PEG (^#^p<0.05). At t = 8 days Plain Synomag-D is significantly different from PEG and PEG-Mannose (*p<0.05).

A drawing of the drainage to lymph nodes in the mouse after a footpad injection is shown in Figure 5A. The first draining node (or SLN) is the pLN, and the HENs include the sciatic, iliac and renal lymph nodes. Figure 5B shows representative 2D images acquired at 4 hours and 8 days post-injection, scaled differently from Figure 4 to highlight HEN signal. MPI signal could be detected in some HENs for all of the Synomag-D particles. In Figure 5C the signal in the HENs is displayed as the percentage of the SLN signal (%SLN) for the 4 hour, 24 hour and 8 day timepoints since HENs were never visible at 10-20 minutes post-injection. Signal is most consistently seen in the sciatic lymph node for all SPIONs, and least consistency seen in the renal lymph node–which is the furthest draining lymph node from the injection site. For mice administered

**Figure 5.**
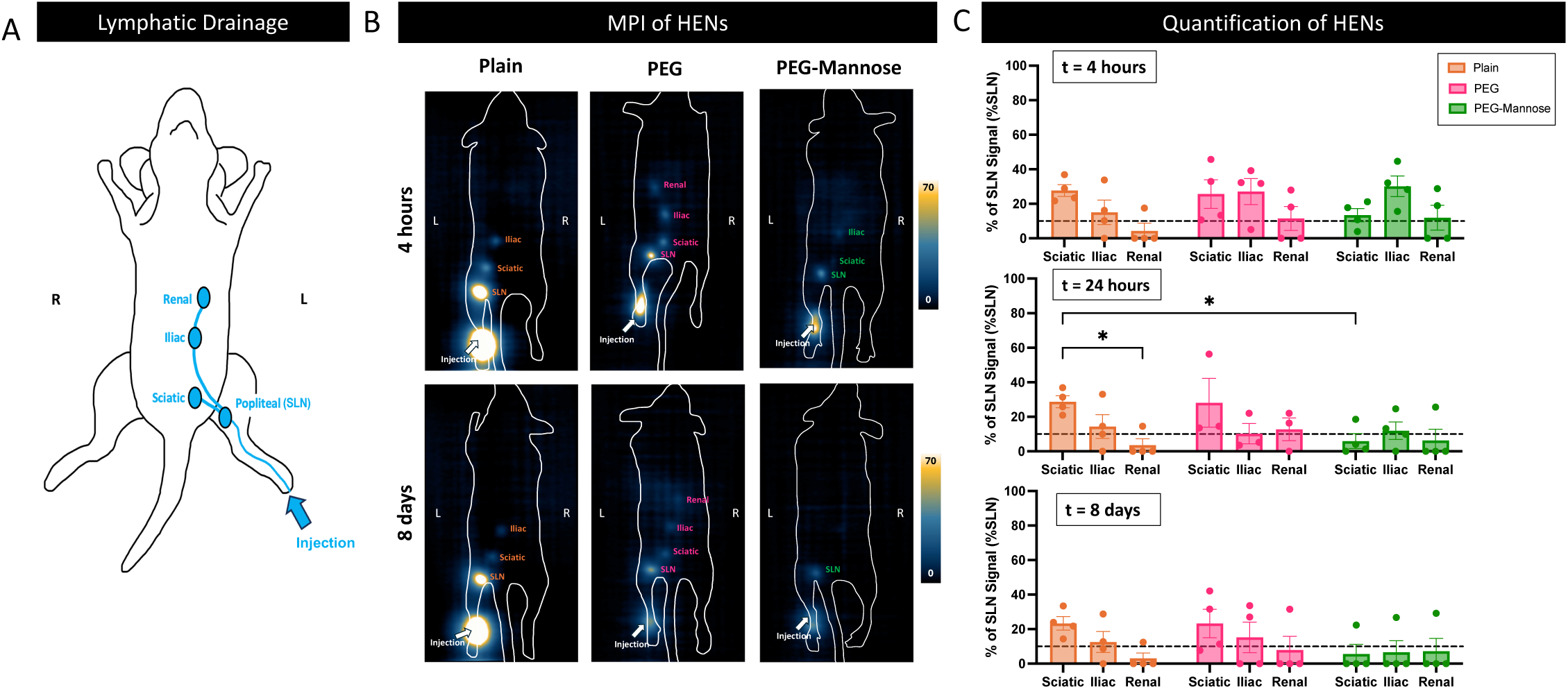
Comparing Signal at the HENs for Synomag-D with Different Surface Coatings. (A) A schematic depicting the lymphatic drainage of mice upon footpad injection is shown. The first draining lymph node in this model is the popliteal lymph node (pLN) which is designated as the SLN. Next SPIONs can subsequently drain to the sciatic, iliac, and renal lymph nodes. (B) Representative MPI is shown for mice administered Synomag-D SPIONs with different surface coatings (Plain, PEG, and PEG-Mannose) 4 hours and 8 days after injection. (C) The %SLN is quantified for the HENs 4 hours, 24 hours, and 8 days post injection (*p<0.05). A dashed line through y=10 is included to represent the 10% rule for determining if echelon nodes should be removed.

Synomag-D Plain and PEG, HEN signal was seen at all timepoints. For the PEG-Mannose-coated Synomag-D, HEN signal was reduced 8 days after injection, with HEN signal seen in only 1 mouse after 8 days. A dashed line is included at y = 10% (Figure 5C) to illustrate the 10% rule for determining if echelon nodes should be removed. Nodes with signal above 10% would be removed during SLNB. At t = 4 hours, based on mean MPI signal, the sciatic and iliac nodes would be removed for all surface coatings. At t = 8 hours, sciatic and iliac nodes would only be removed for the Plain-and PEG-coated Synomag-D.

### Ex vivo MPI of SLNs

Ex vivo imaging was performed on fixed nodes after the last imaging time point (Figure 6A) and signal was detected for all SPIONs except VivoTrax+. The total MPI signal measured in vivo on day 8 is compared to the signal measured ex vivo with the same imaging parameters (Figure 6B). There was no significant difference in MPI signal between ex vivo and in vivo signal for any of the tested SPIONs, validating in vivo results.

**Figure 6.**
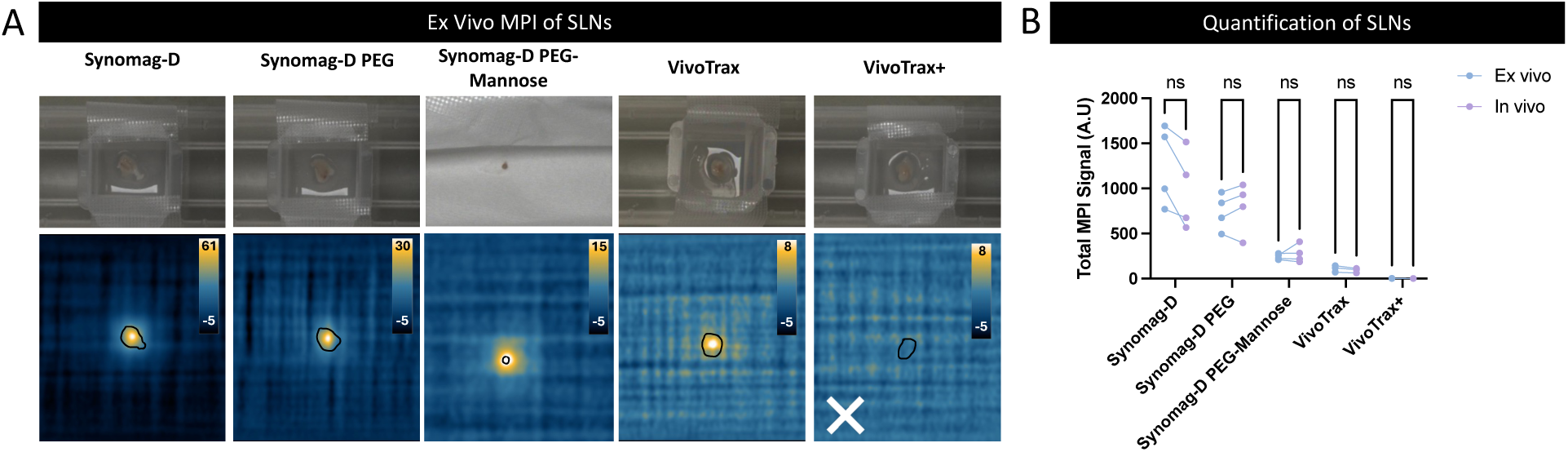
Ex vivo MPI of SLNs. (A) Representative ex vivo MPI of SLNs for each SPION. For VivoTrax+, the white “x” indicated that signal was not detected for any mice ex vivo. (B) Ex vivo MPI signal was quantified and related to day 8 in vivo signal. There is no significant difference between in vivo (day 8) and ex vivo total MPI signal for any of the SPIONs.

### Histology of SLNs and HENs

Lymph nodes removed after the final imaging session were stained and qualitatively assessed using PPB staining (Figure 7A). Examples of lymph node dissections are shown in **Figure S4**. Iron staining was present for all nodes that showed MPI signal in vivo. No staining was seen in nodes obtained from control mice that were not injected with SPION. Most iron staining was seen in the lymph node sinuses. Qualitatively it appeared that there was more iron staining in the Synomag-D nodes compared to the VivoTrax/VivoTrax+ nodes. Additionally, for the Synomag-D particles, iron can also be seen in the trabeculae and closer to the efferent lymph vessel. This is more obvious for Synomag-D PEG, where iron can be seen in the medullary sinus. This suggests that these particles flow towards the efferent vessel and the HENs, consistent with our in vivo findings.

**Figure 7.**
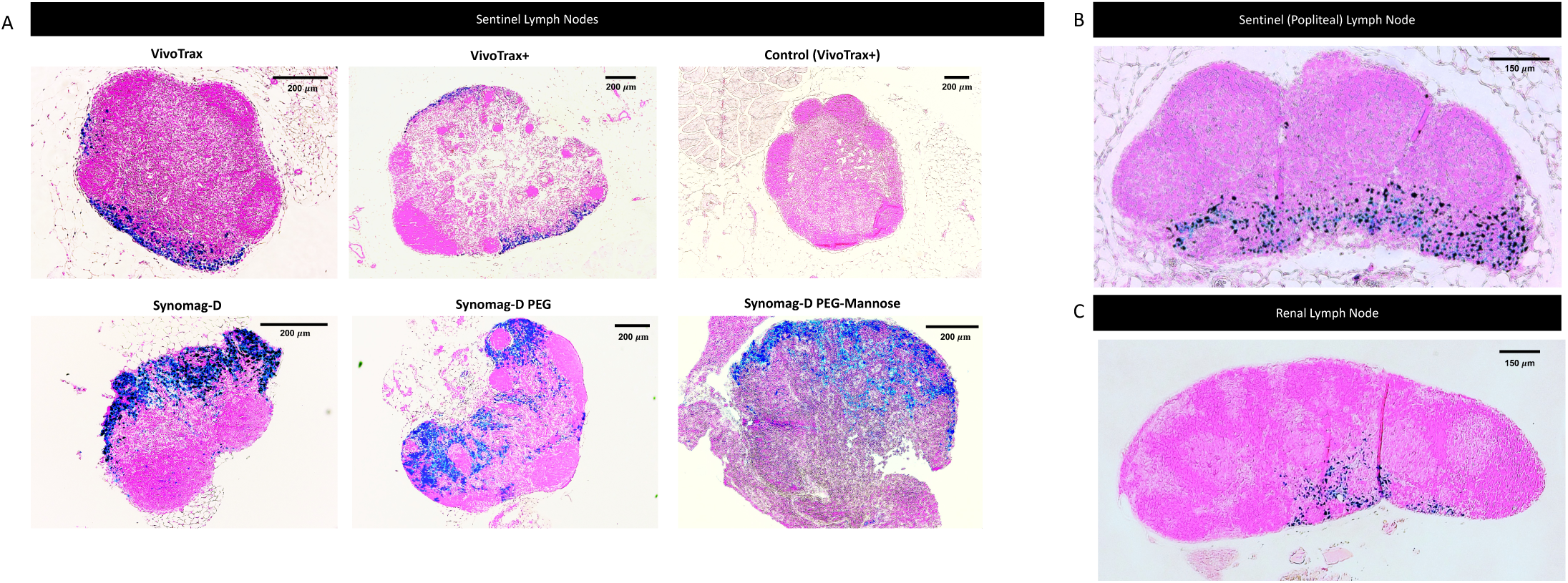
Histology of SLNs and HENs. (A) Perl’s Prussian Blue staining (iron = blue) of a representative SLN from each SPION. Scale bars are on the top right of each image. (B) and (C) show a SLN and a HEN from a mouse administered Synomag-D PEG.

For a mouse administered Synomag-D PEG, iron staining is shown for the SLN (Figure 7B) and the renal lymph node (Figure 7C), which is the last HEN where MPI signal was seen. Qualitatively there was less iron staining in the renal lymph node compared to the SLN in this mouse, which confirms what was quantified from imaging.

### ICP-MS

ICP-MS provides quantification of various elements. Here we quantified iron in the ipsilateral and contralateral (control) SLN for one mouse from each SPION and compared it to the estimated iron from MPI quantification.

## Discussion

In this study we show that the magnetic properties and lymphatic pharmacokinetics of SPIONs are both important factors for identifying draining lymph nodes with MPIL. The MPI signal in the draining SLN was first compared after the administration of three particles: VivoTrax, VivoTrax+ and Synomag-D. In full FOV images of mice the MPI signal was significantly higher for Synomag-D compared to VivoTrax. The %ID measured in the SLN for Synomag-D was 2-3x higher than VivoTrax at the 4 and 24 hour imaging timepoints and 6x higher at 8 days. The particle sensitivity, assessed in vitro with MPR, showed that the peak MPI signal for Synomag-D was 2.5x higher than VivoTrax. This suggests that the accumulation of these two SPIONs in the SLN may be similar at the earlier timepoints and that the different magnetic properties could be responsible for the differences in MPI signal. MPI signal was visible in distal nodes for mice injected with Synomag-D but not VivoTrax.

In theory the ideal MPI tracer should be monodisperse and have a magnetic core diameter of approximately 20-25 nm.^34^ Although widely used for MPI, VivoTrax is not considered optimal due to its bimodal size distribution. It has been reported that ∼70% of the particles are single 5 nm iron cores and ∼30% of the particles are aggregates of these cores (multicore particles) which measure between 20-30 nm.^35,36^ The smaller cores do not magnetize sufficiently, leaving only the small fraction of multicore particles that contribute to the MPI signal. Synomag-D was synthesized specifically for MPI applications. The particles are described as nanoflowers because of the appearance of the clusters of iron cores by electron microscopy. The average size of the nanoflowers are approximately 30 nm and studies have reported between 2.5 and 4x peak MPI signal compared to Resovist or VivoTrax.^25,30,37,38^ The greater difference in %ID at the last timepoint may be related to higher retention of Synomag-D compared to VivoTrax. This is supported by histology of the SLNs removed at 8 days which showed more iron staining for Synomag-D compared to VivoTrax. In addition, only 3 of the 4 mice injected with VivoTrax still had SLN signal at 8 days post-injection.

No MPI signal was detected in full FOV images of the SLN after administration of VivoTrax+. This was not expected considering that MPR shows that the peak MPI signal for VivoTrax+ is significantly higher than VivoTrax. VivoTrax+ is a magnetically fractionated form of VivoTrax that selects for the fraction of larger cores for improved MPI performance, which explains the higher particle sensitivity compared to VivoTrax as measured by MPR. This suggests that the low in vivo sensitivity for VivoTrax+ is related to reduced lymphatic drainage to the SLN.

While the iron core size is critical to the intensity of the MPI signal, the total hydrodynamic size of SPIONs (core size plus coating) and the particle coating affect the biological fate and trafficking to lymph nodes. Nanoparticle delivery to lymph nodes has been an active area of research for many years. After intradermal or subcutaneous administration, nanoparticles traffic to lymph nodes via interstitial transport, or they are transported by dendritic cells. Though studies do not identify an exact optimal size, results suggest that the hydrodynamic size for the most efficient interstitial transport is between 20 and 50 nm and that larger particles only later reach the LNs by dendritic cell transport. Mori et al. examined particles with hydrodynamic sizes ranging from 50 nm to 1,000 nm and found that those 200 nm or larger were unsuitable due to minimal uptake within 24 hours. Iida et al. evaluated citrate-coated nanoparticles of 4 nm, 8 nm, and 20 nm, reporting that the 20 nm particles were retained most effectively in the SLNs and reached the highest concentration. The manufacturer reported hydrodynamic diameters for Synomag-D, VivoTrax, and VivoTrax+ are 70, 62, and 62 nm respectively. Our DLS measurements for VivoTrax and VivoTrax+ gave hydrodynamic diameters of 57.48 nm and 91.06 nm. The larger diameter for VivoTrax+ may play a role in its inferior drainage to the SLN.

Full FOV images of mice were used to guide the subsequent acquisition of small FOV images focused on the SLN (Figure 3). These images have a higher signal-to-noise ratio because a small FOV limits the spatial domain over which noise contributes and excludes off-target background signal, such as the injection site which has much higher tracer concentration compared to the SLN. In these images MPI signal was detected in all SLNs for all three SPIONs. The higher sensitivity afforded by imaging a small focused FOV revealed that there was some accumulation of VivoTrax+ in the SLN. Although the signal for VivoTrax+ was still lower than that for VivoTrax there was no statistically significant difference at any imaging timepoint. For clinical MPI whole body imaging is not practical.^39,40^ The first human sized MPI scanners have been designed for imaging specific regions of the body, for example the torso, brain and thigh.^41–44^ It is more realistic to imagine acquiring a full FOV of a particular body region and using it to guide the prescription of a small FOV focused on a lymph node basin. Scaling MPI technology to human sizes will lead to reduced sensitivity compared to pre-clinical MPI scanners.^45^ The gain in sensitivity with a small focused FOV may help to offset this. 3D imaging can also be used to increase MPI sensitivity. **Figure S5** shows 3D whole mouse body images acquired at 4 hours post-injection for each of the SPIONs tested. The SLNs could be detected in 3D images for all SPIONs. In some cases, HENs that were not detectable in 2D images could be visualized with 3D MPI. However, this improvement in detection sensitivity comes with a significant time penalty; the 3D full FOV image acquisition took 35 minutes compared to 2.5 minutes for 2D MPI.

While the effect of nanoparticle size on lymphatic transport is well established there is less data about the optimal nanoparticle surface chemistry required for maximizing lymph node accumulation of nanoparticles after intradermal administration. To study the influence of the surface coating we compared three types of Synomag-D particles with similar hydrodynamic and core sizes. MPI signal was detected at the SLN in all mice for all three Synomag-D particles, at all timepoints except for in the earliest images. At 10 minutes after the particle administration, MPI signal was detected at the SLN in all mice administered the Plain Synomag-D, in one mouse administered PEG-coated Synomag-D, and in none of the mice administered the PEG-Mannose-coated Synomag-D. Particle sensitivity measured by MPR showed that the peak MPI signal for the Plain Synomag-D was significantly higher than Synomag-D PEG and Synomag-D PEG-Mannose. This may explain our observations at the earliest imaging timepoint. At all other timepoints the %ID was highest for Synomag-D PEG, intermediate for the Plain Synomag-D, and lowest for Synomag-D PEG-Mannose, although there were no statistically significant differences (Figure 4).

PEGylating nanoparticles is a well-known strategy to enhance interactions with biological materials and extend the blood circulation time.^46,47^ Several early studies demonstrated that coating nanoparticles with PEG can also enhance lymph node accumulation after intradermal or subcutaneous administration in mice.^48,49^ McCright et al., found that PEGylation of polystyrene nanoparticles improved transport across lymphatic endothelial cells and into lymphatic vessels increasing the number of particles that accumulate in lymph nodes, compared to unmodified particles.^50^ Our results are in agreement with these studies, although we expected to observe a more substantial difference in the MPI signal at the SLN for Synomag-D PEG compared to Plain Synomag-D. It is possible that the transport of Synomag-D PEG to the SLN is greatly improved over the Plain Synomag-D but that this was not reflected in the images because of the inferior particle sensitivity, as measured by MPR.

The use of surface conjugated ligands is another strategy for targeting specific lymphatic cells to enhance accumulation of nanoparticles in lymph nodes. Mannose is a ligand for targeting CD206 receptors on macrophages and dendritic cells. Technetium (^99m^Tc) tilmanocept (Lymphoseek) is a mannose containing radiolabeled tracer used for lymphatic mapping which shows rapid clearance from the injection site, high retention in the SLN, and low distal node accumulation, compared to non-targeted radiocolloids.^7^ The rapid clearance and accumulation in nodes is related to its small size (7 nm compared to ∼100 nm for conventional radiocolloids). The high retention in nodes and low flow through is attributed to specific binding to nodal reticuloendothelial cells. In HNC where lymphatic drainage pathways are complex the use of non-targeted radiocolloids have well known limitations, including injection site shine-through which masks SLN signal and rapid flow through to HENs, which both result in the risk of false negatives. A meta-analysis by Rovera et al., looked at 24 studies comparing non-targeted radiocolloids (^99m^Tc-sulfur colloid) to ^99m^Tc-tilmanocept for breast cancer, melanoma, and HNC.^51^ For breast cancer, melanoma, and HNC, ^99m^Tc-tilamanocept and ^99m^Tc-sulfur colloid showed comparable detection of at least one SLN, and quick clearance from the injection site. Some studies have also shown that for breast cancer and melanoma ^99m^Tc-tilamanocept allowed for removal of fewer lymph nodes, which may be attributed to its specificity for CD206 receptors.^52,53^ For HNC, benefits of ^99m^Tc-tilamanocept over non-targeted radiocolloids for the number of nodes removed is not described well. The rapid clearance of ^99m^Tc-tilamanocept may be particularly advantageous for floor of mouth tumours where shine-through from the injection site can impede signal in nearby nodes.^54^

Krishnan et al. evaluated a mannose-labeled SPION called Ferrotrace (74 nm), with respect to SLN uptake and distribution to HENs, in a large animal model of HNC.^55^ MRI was performed at 15-30 minutes, 6 hours, 7 days and 26 days post-injection to visualize the signal loss caused by iron in lymph nodes. The amount of iron in nodes identified by MRI was measured by scanning them using a magnetometer probe. MRI showed rapid flow of tracer to three lymph nodes, the SLN and a second and third echelon node. Ferrotrace continued to be taken up into nodes over 4 weeks in a linear fashion. The %ID for SLNs was between 6 and 22%. There was significantly higher uptake in SLNs compared to HENs; on average 83% of the iron resided in the SLNs compared to 15% for the second echelon and 6% for the third echelon nodes, suggesting high specificity of Ferrotrace for the SLN. Histology of the nodes showed that the SPION particles were found mostly in the cytoplasm of macrophages.

We compared a mannose labeled Synomag-D PEG tracer with Plain Synomag-D and Synomag-D PEG. MPI showed signal in four lymph nodes, the SLN (popliteal lymph node) and three HENs (sciatic, iliac and renal). There was no statistically significant difference in the MPI signal at the SLN for the three tracers at timepoints between 4 hours and 8 days post-injection. This agrees with findings from studies which compared ^99m^Tc-tilamanocept and non-targeted radiocolloids which report comparable detection of SLNs.^56^ The %ID for SLNs was between 7 and 31%, similar to the %ID measured by MRI in the Krishnan study. There was significantly higher uptake of iron in SLNs compared to HENs in Synomag-D PEG-Mannose administered mice which is also similar to the findings in the Krishnan paper for the mannose-labeled SPION Ferrotrace. For nodes that were detectable, on average 79% of iron resided in the SLN, 5% in the sciatic lymph node, 10% in the iliac lymph node, and 10% in the renal lymph node. We saw significantly lower signal at the injection site (higher clearance) for the PEG and PEG-Mannose administered mice compared to the Plain Synomag-D, with the highest signal drop-off 8 days post-injection. There was no significant difference between PEG and PEG-Mannose, with both reaching an injection site clearance half time at ∼ 8 days. Future work would be required to determine how long signal persists in the injection site and how quickly signal fully drops off.

MPI signal was only detected in HENs after administration of the three Synomag D SPIONs (Figure 5). In most cases, for HENs that were detectable, the %SLN did not change markedly between 4 hours and 8 days post-injection. For Plain Synomag-D and Synomag-D PEG the MPI signal was usually highest in the second echelon node (sciatic), lower in the third echelon node (iliac) and lowest in the fourth echelon node (renal). Overall, flow through to HENs was lowest for Synomag-D PEG-Mannose, with only one out of four mice having detectable signal in all three HENs after 8 days.

## Limitations

This study used naïve, non-tumour-bearing mice. The first draining lymph node from the intradermal footpad injection, the popliteal node, was referred to as the SLN. This model allowed for a straightforward analysis of the lymphatic drainage of different SPIONs. Tumours alter local lymphatic drainage in several ways including physically blocking lymphatic vessels and nodes, increasing interstitial pressure, and stimulating the formation of new lymphatic vessels.^57,58^ Future studies using metastatic and non-metastatic tumour models in mice will be important to further evaluate MPIL. Clinically, the echelon nodes are defined as those that drain from the SLN in the same basin. In this study lymph nodes distant to the popliteal node represented higher echelon nodes (HENs) and allowed us to learn about the potential for flow through of the different SPIONs.

The effect of different doses of iron was not thoroughly explored in this study. The dose used in mice was scaled by weight from the dose approved for humans (56 mg of Fe/injection). A single mouse was imaged after a half dose of Synomag-D was injected. **Figure S6** shows that the MPI signal in HENs can be reduced by reducing the amount of iron that is injected; only the second echelon node (the sciatic) was detected and only up to 24 hours post-injection. Despite the lower dose, strong signal was still detected in the SLN (%ID ∼ 15%) over the course of 8 days, showing the potential for testing even lower doses.

Our histological analysis of node tissues was limited to a qualitative assessment of PPB staining of node sections which only provided information on the presence and general location of iron particles. More detailed microscopy is required to better understand the total accumulation of SPIONs in nodes. Immunohistochemistry to identify specific immune cell types in nodes could be compared to PPB staining to determine whether iron particles are bound to or internalized in CD206 dendritic cells or macrophages and if this is more prevalent for Synomag-D PEG-Mannose.

## Conclusions

In this study, we demonstrated the feasibility of MPI to identify SLNs labeled by SPIONs in a mouse model. Magnetically labeled draining lymph nodes were detected and monitored by MPI for 8 days and then excised to verify iron presence by ex vivo MPI and histology. The pharmacokinetics of SPIONs with different magnetic properties and surface coatings was investigated to determine signal strength at the SLN and flow through to distant nodes. SPIONs for SLNB should have rapid drainage, high SLN accumulation and high specificity to the SLN. This study shows that for MPIL high particle sensitivity is also important.

Our results contribute to the growing body of evidence that show magnetic SLN mapping is a viable alternative to conventional methods. MPI could play a complementary role by providing the surgeon with pre-operative images showing the location of the SLN(s) for sensitive and specific SLNB while the magnetometer provides numerical counts to guide the surgeon intra-operatively. Further studies will determine if MPI could be used alone for both sensitive detection and quantitative imaging of the SLN.

## Acknowledgements

We would like to acknowledge the IMPAKT Facility at Western University, and funding from the Canadian Institutes of Health Research and the London Regional Cancer Program.

## Competing Interests

Dr. Benjamin Fellows was employed by Magnetic Insight Inc. Dr. Olivia Sehl is employed Magnetic Insight Inc.

## Supplementary Figures

**Figure S1.**
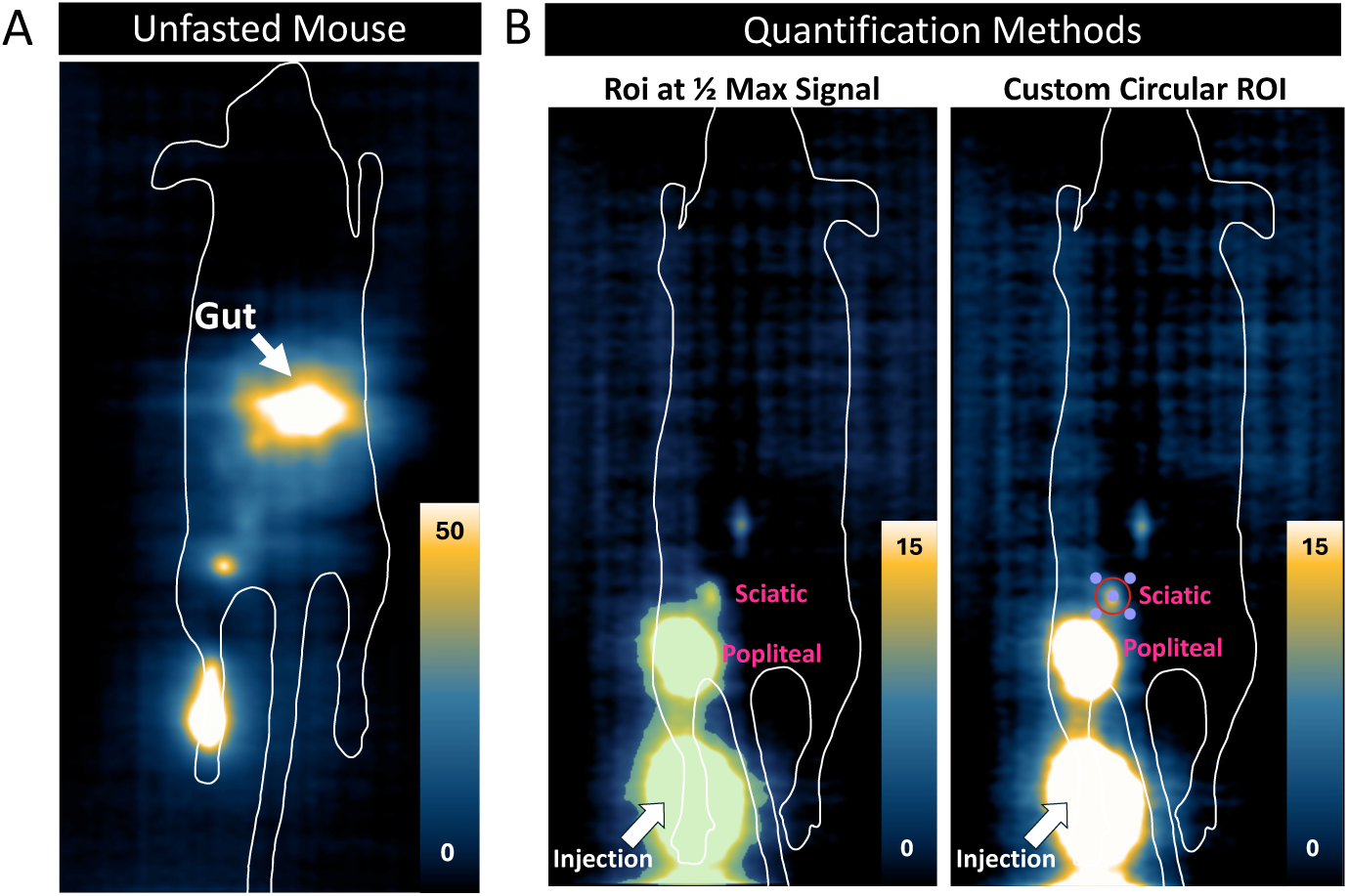
Considerations when imaging and quantifying mouse lymph nodes. (A) An unfasted mouse administered Synomag-D PEG 24 hours after injection is shown. Signal in the gut due to iron in mouse feed impedes on imaging of HENs. (B) There must be careful selection of quantification methods. A mouse administered Synomag-D PEG is shown 4 hours after injection. If an ROI at ½ the maximum signal of the sciatic lymph node is applied, we are unable to isolate sciatic lymph node from the SLN and injection site. In this case, accurate quantification of the sciatic lymph node is not possible. Instead, a custom circular ROI of the same size is drawn around all lymph nodes, allowing for signal isolation.

**Figure S2.**
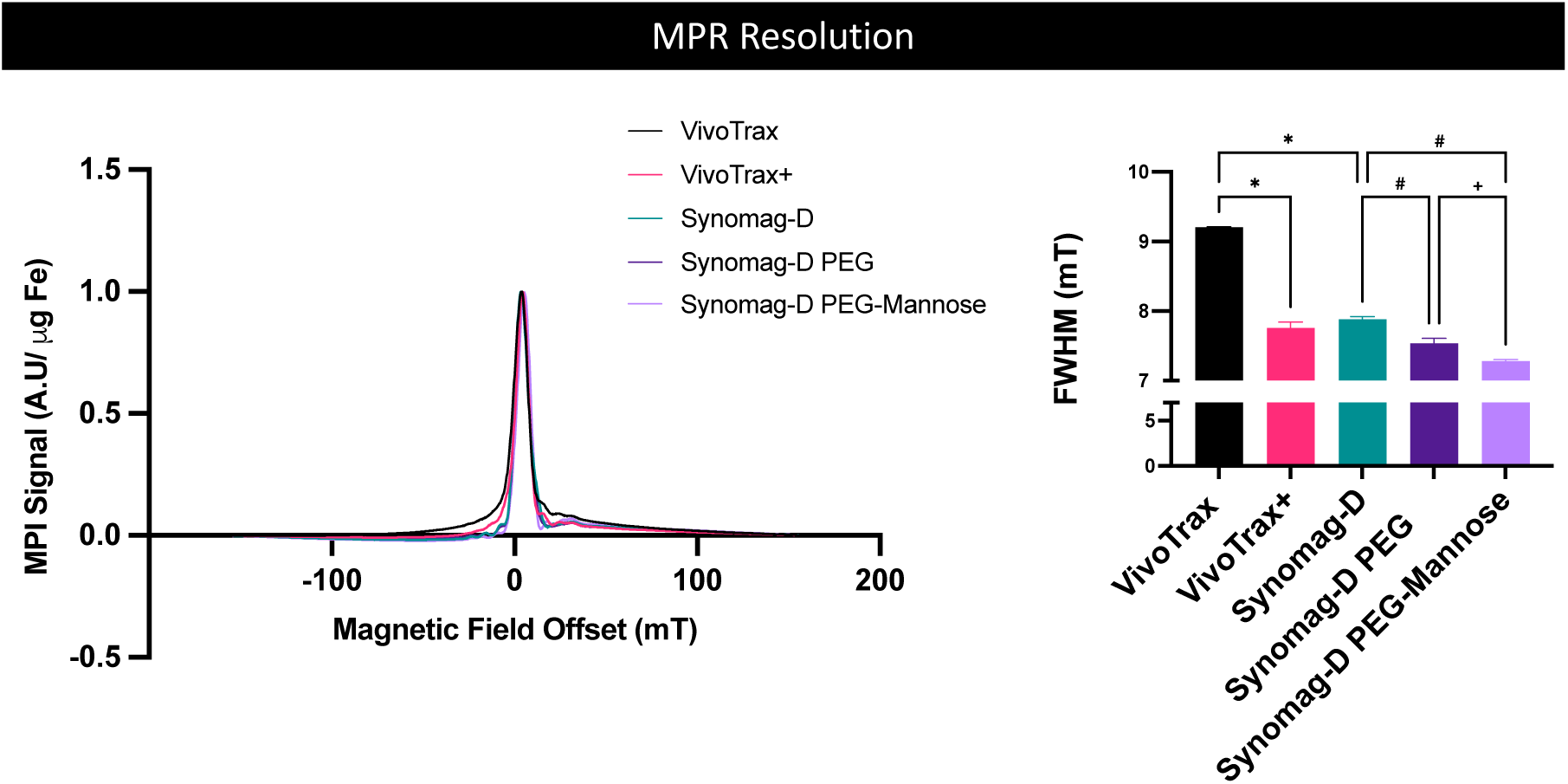
MPR Resolution. Particle resolution, measured by FWHM is shown. The bar graph compares FWHM for each SPION (n=3 for each SPION), where lower values indicate superior resolution (*p<0.05 (paired t-test with VivoTrax), ^#^p<0.05 (paired t-test with Synomag-D), ^+^p<0.05 (paired test with Synomag-D PEG)).

**Figure S3.**
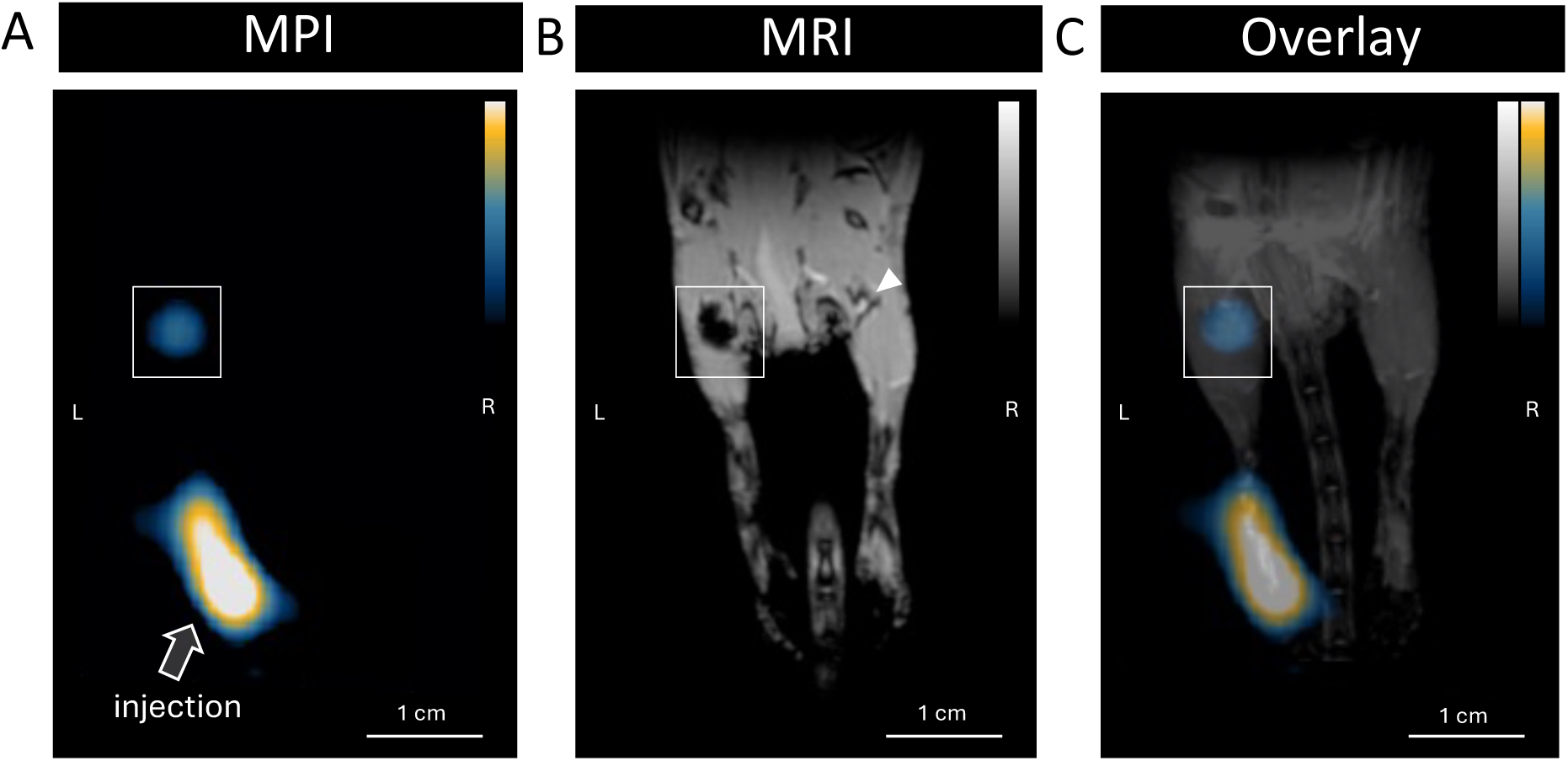
Dual modality imaging of SLNs. For one mouse administered VivoTrax, both MRI and MPI scans were obtained 4 hours post-injection. (A) A MPI scan is shown, with a white box around the SPION signal in the SLN. An arrow is pointing to the injection site. (B) The corresponding MRI scan provides anatomical information and the SPIONs at the injection site and the SLN can be seen as signal voids. (C) An overlay of the MPI and MRI scans is shown.

**Figure S4.**
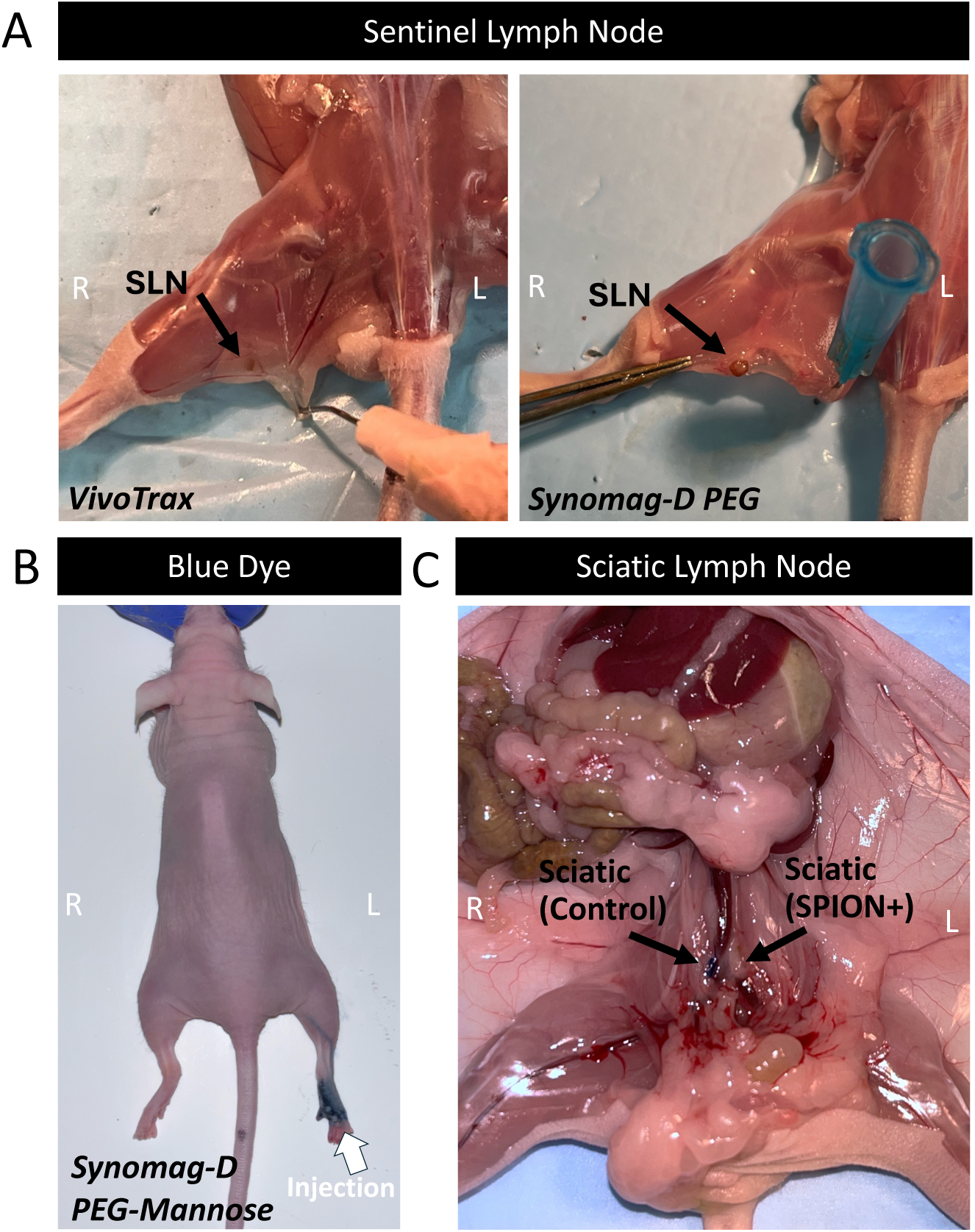
Dissection of SLNs and HENs. (A) Iron laden SLNs for mice administered VivoTrax and Synomag-D PEG are shown. The lymph nodes are visibly brown. (B) When performing dissections, 25 uL of 5% Evans Blue Dye was injected into contralateral footpad to guide dissections. (C) The contralateral (control) sciatic lymph node and ipsilateral (SPION+) sciatic lymph node are shown.

**Figure S5.**
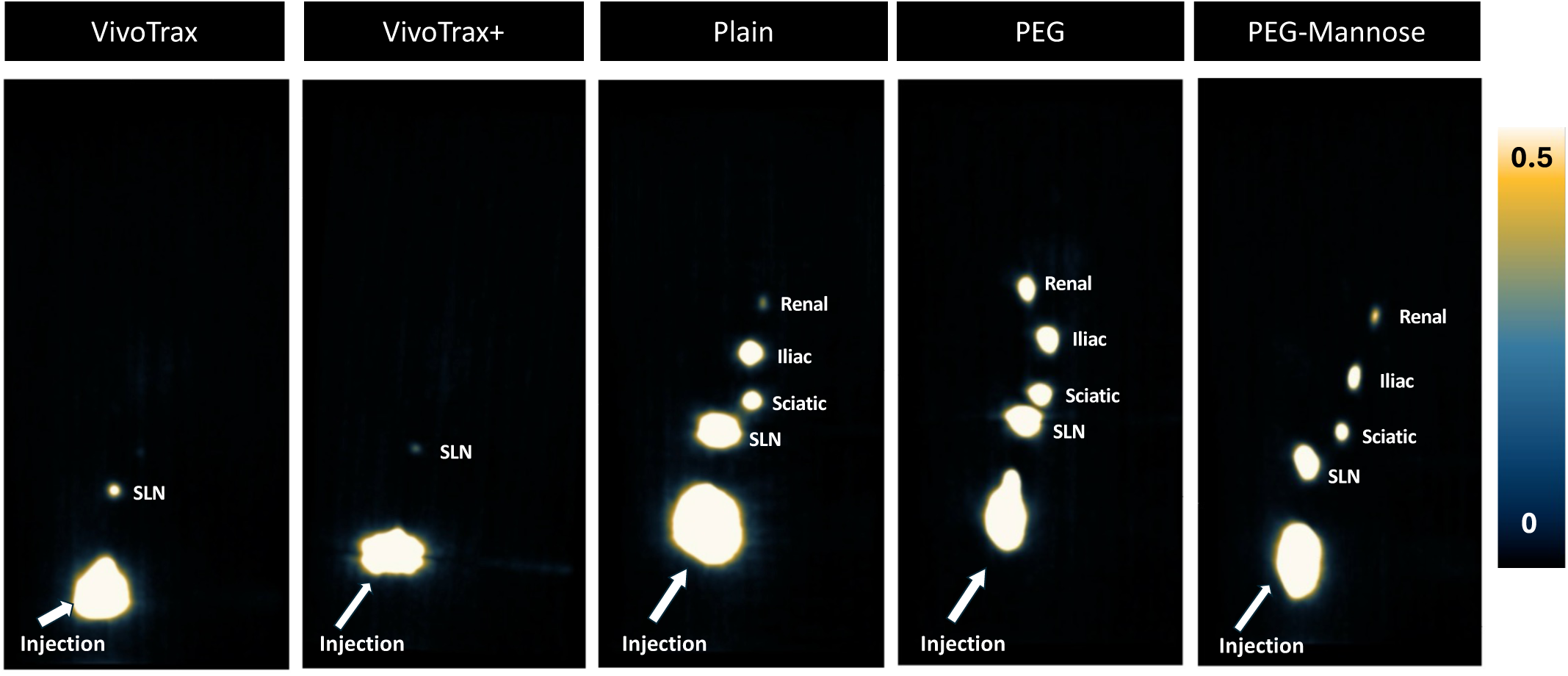
3D MPI of each SPION. 3D MPI was acquired for once mouse from each SPION at the t = 4 hour timepoint. The SLN can be detected for all mice. Additionally, all HENs (sciatic, iliac, and renal) can be seen for the different for Synomag-D surface coatings.

**Figure S6.**
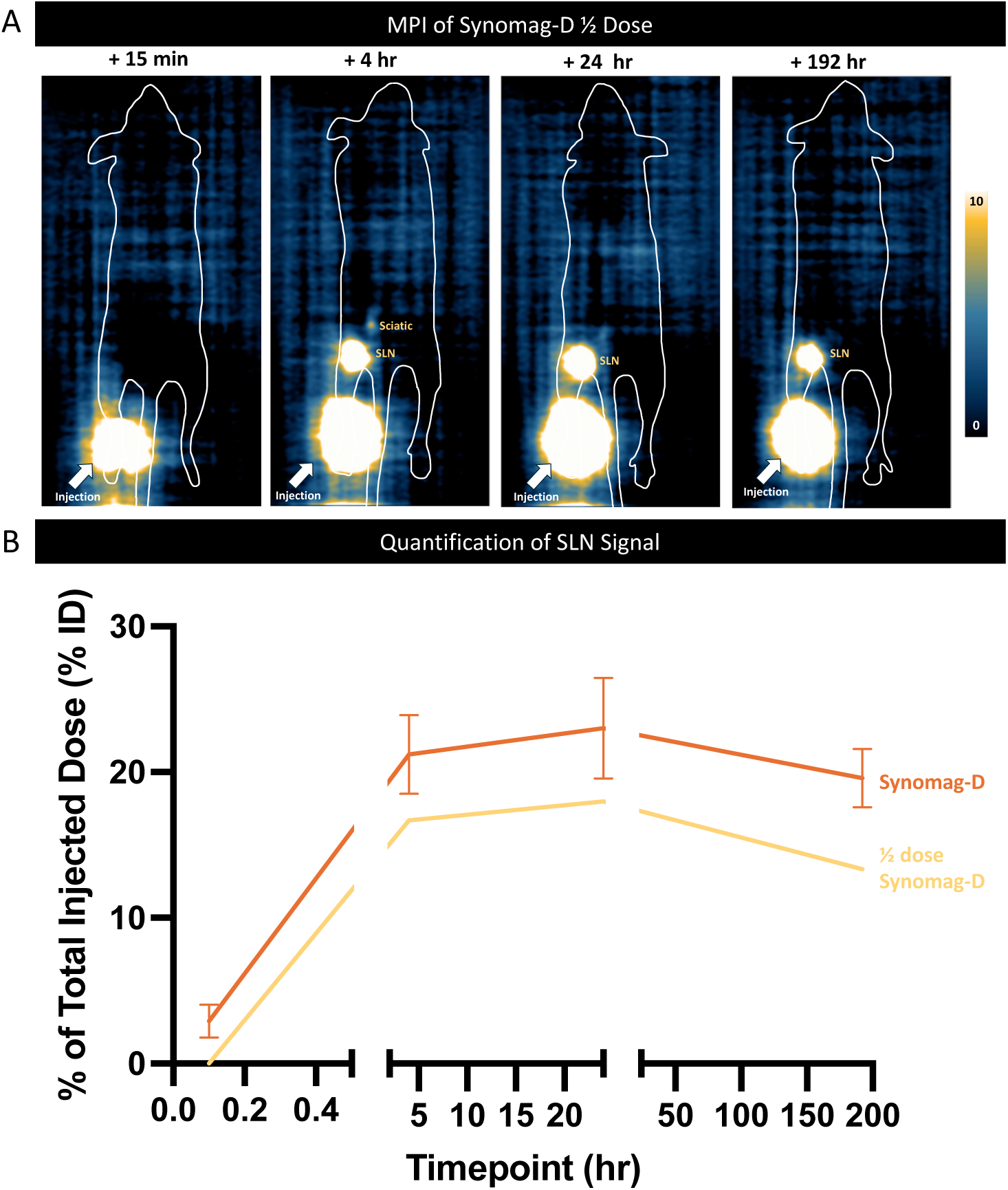
Administering half the tested dose of Synomag-D. (A) One mouse was administered ½ a dose (0.4 mg/kg) of Synomag-D (Plain) and imaged 10-20 minutes, 4 hours, 24 hours, 192 hours (8 days) post-injection. MPI is shown at all timepoints. The ½ dose reduced uptake to HENs. The sciatic lymph node was only detected 4 hours post-injection. (B) Quantification of SLN signal for the ½ dose (n = 1) is compared to mice administered the full dose of Synomag-D (n = 4).

